# A quantitative eDNA-based method to monitor fish spawning in lakes: application to European perch and whitefish

**DOI:** 10.1101/2022.05.05.490723

**Authors:** Marine Vautier, Cécile Chardon, Chloé Goulon, Jean Guillard, Isabelle Domaizon

## Abstract

There is an urgent need to evaluate the effects of anthropogenic pressures and climatic change on fish populations’ dynamics. When monitored in lakes, the spawning of fish is generally assessed using traditional, mostly destructive or damaging, methods as gillnetting and collection of fertilized eggs.

Over the last decade, environmental DNA (eDNA) based methods have been widely developed for the detection of aquatic species, offering a non-invasive alternative method to conventional biomonitoring tools. In particular, the emergence of new methods as the droplet digital PCR (ddPCR) offer the possibility to quantify an absolute eDNA signal in a very sensitive way and at a low cost.

Here, we developed and implemented a quantitative eDNA method to monitor the spawning activity of two fish species, European perch and whitefish. ddPCR protocols were formalized based on existing and newly designed COI primers, and were applied during four spawning periods in lake Geneva.

The results demonstrate the efficiency of eDNA coupled with ddPCR to identify the timing and duration of the spawning periods, as well as the peak of the spawning activity for the targeted species. In addition, the use of a control species (i.e., quantification of the eDNA signal of a fish that does not reproduce during the monitoring period) was shown to be relevant to clearly discriminate fluctuations of the eDNA signal associated to the spawning activity from the baseline eDNA signal. For future implementation, we recommend using an integrative sampling strategy (e.g., pooled samples for a give station) to smooth the local variability of the eDNA signal. These results show that we reached an operational level to use these non-invasive eDNA methods to monitor the spawning periods of these two fish species in large lakes.

## Introduction

Environmental DNA (eDNA) methods are increasingly used as monitoring tool for freshwater organisms (Pawlowski *et al*. 2020). Over the last decade, the use of eDNA has been widely developed for the detection of species, populations or biological communities (e.g. Goldberg et al. 2011; Jerde et al. 2011; Flynn et al. 2015; Valentini et al. 2016). Consequently, eDNA surveillance is now considered a valuable tool to screen for the presence of aquatic organisms, including rare species (Rees et al. 2014), with an excellent potential for applications to lakes and rivers biomonitoring (e.g. Valentini *et al*. 2016, Lefrancois *et al*. 2018). This non-invasive method offers many advantages over traditional techniques, including ease of species identification, lower sampling costs, and detection of low signals (Ficetola *et al*. 2008; Taberlet *et al*. 2012; Thomsen *et al*. 2012; Bohmann *et al*. 2014). More specifically, the application of eDNA methods for inventorying fish species has benefited from various methodological improvements (Bessey et al 2019; Miya *et al*. 2020) to offer an alternative to conventional surveying tools. Recent applications also shown the potential of eDNA to monitor fish spawning periods (Thalinger *et al*. 2019; Bylemans *et al*. 2017), which is crucial for fish species conservation policy and management.

To monitor fish spawning activity, methods are generally invasive (e.g. gillnetting and collection of fertilized eggs) or injurious (e.g. electric fishing, acoustic telemetry), and the non-invasive methods (e.g. visual surveys and acoustic surveys) are sensitive to observer biases and taxonomic misidentifications (Lefort *et al*. 2015; Bylemans *et al*. 2017). eDNA methods could represent an alternative approach if the applied eDNA protocols are relevant to assess the biomass and/or abundance of fish species precisely enough. Correlation between fish eDNA concentration in water and its spawning activity has been shown in rivers for diadromous (Bergman *et al*., 2016; Gingera *et al*., 2016; Tillotson *et al*., 2018; Antognazza *et al*., 2019; Bracken *et al*., 2019) and potamodromous fish species (Thalinger *et al*., 2019). For non-migratory fish species, different DNA-based approaches have been developed in rivers, as the survey of the spawning activity of the Australian Macquarie perch *(Macquaria australasica)*, by comparing ratios of nuclear and mitochondrial eDNA (Bylemans *et al*., 2017) or the correlation between eDNA signal and the probability to observe bigheaded carps (*Hypophthalmichthys spp*.) ichthyoplankton (Fritts *et al*., 2019). Though some correlations have been established between eDNA signal (in water) and fish abundance in lakes (*e*.*g*., Nevers *et al*., 2018; Klobucar *et al*., 2017), only one eDNA metabarcoding approach has been performed to monitor Artic char (Salvelinus alpinus L.) spawning periods in lakes (Muri et al., 2022).

Anthropogenic pressures occurring at local scale (*e*.*g*., artificialization of the shoreline, eutrophication) and at global scale (e.g. climatic warming) have increased the interest in studying the impact of environmental changes on the key stages of the fish life cycle. Those pressures affect fish reproductive success and thus fish abundance, as reported for instance for peri-alpine lakes subjected to changes in phosphorus concentration (eutrophication and reoligotrophication) and modifications of thermal conditions (Massol *et al*. 2007). In particular, increased water temperatures and shortened winters may result in the modification of both the timing of spawning and the duration of eggs incubation (e.g. Stewart *et al*., 2021; Lyons *et al*. 2015 ; Starzynski and Lauer, 2015). In Lake Geneva, climate change and re-oligotrophication are known to affect percid and salmonid populations whose stocks are subject to significant fluctuations (Gillet *et al*., 2013; Caudron *et al*., 2014; Mari *et al*., 2016), with societal consequences on the fishing activity (Nõges *et al*., 2018). To understand these fluctuations, the monitoring of the spawning phenology of two fish species which are important for recreational and commercial fisheries, *i*.*e*., perch (*Perca fluviatilis*) and whitefish (*Coregonus lavaretus*), has been set up in lake Geneva. For perch the traditional monitoring is traditionally performed using artificial spawning substrates placed at a reference site to collect egg strands (from early April to late June) (Gillet *et al*., 2013), while for whitefish, the traditional monitoring is based on the use of gillnets deployed each week (from late November to late January) in an area where whitefish naturally spawns (Goulon *et al*., 2021). Recent advances in the quantification of eDNA signals from water samples (e.g. Nathan *et al*., 2014; Doi *et al*., 2015) open up high potential alternatives to circumvent or efficiently complement these traditional and invasive methods.

The aim of this study was therefore to test our capacity to monitor the timing of the perch and whitefish reproduction periods using a quantitative eDNA method, assuming that, when spawning, fish gather on spawning areas, have an increased activity and release gametes, leading to higher DNA amounts released in the surrounding environment. Water was collected in the littoral zone, where perch and whitefish spawn, and we tested two different sampling strategies : a replicated discrete sampling and an integrated sampling strategy to smooth the potential local heterogeneity of the eDNA signal (Sato *et al*., 2017a). To distinguish the basal eDNA signal emitted by a fish species that lives permanently in the lake from the signal revealing its spawning activity, we chose to systematically quantify the eDNA of the two species, one species serving as an “eDNA control” for the other. To reach a high level of sensitivity and detect small variations in the eDNA quantity, we used a recent technology, the droplet digital PCR (ddPCR). The digital PCR (dPCR) approach has the advantage of being quantitative (Nathan *et al*., 2014) and more efficient than qPCR to detect rare aquatic organisms, including fish (Wood *et al*., 2019; Brys *et al*., 2020; Doi, Uchii, *et al*., 2015). Both primers sets used, *i*.*e*., the one developed in this study for perch and the one proposed by Hulley *et al*. (2019) for whitefish, target the cytochrome c oxidase I (COI) mitochondrial gene, and their compatibility with ddPCR analysis was validated following appropriate specificity and sensitivity tests. The results obtained from eDNA quantification were compared with the traditional monitoring carried out for these two species in Lake Geneva.

## Materiel & methods

### Study site and selection of sampling locations (Fig. 1)

Lake Geneva is located in the northern Alps (46°27’N, 06°32’E) at an elevation of 372m. The lake surface area is 580.03 km^2^, with a maximal depth of 309 m. Perch and whitefish are the two most abundant fish species in Lake Geneva, and their abundance strongly varied over the past (Anneville *et al*., 2017). The monitoring of their spawning phenology allowed to estimate the start, the end and the peak of the spawning period for these two species, in order to guide the stakeholders with recommendations to adapt the fisheries closing dates, but also to observe possible modifications from one year to another as for instance the number of spawners, the spawning timing or duration (Goulon *et al*., 2021). In Lake Geneva, the whitefish spawning takes place from early December to mid- January in the littoral zone (Goulon *et al*., 2021). While whitefish live most of the year in the pelagic zone, during the spawning period they daily migrate to the littoral zone at nightfall for the reproduction and then return at deeper depth (pelagic zone) at sunrise. In winter, perch usually school on depth around 30-50 m, and migrate in spring in the shallow littoral zone to spawn. Their spawning period occurs from late April to early June (Gillet *et al*., 2013).

**Figure 1.**
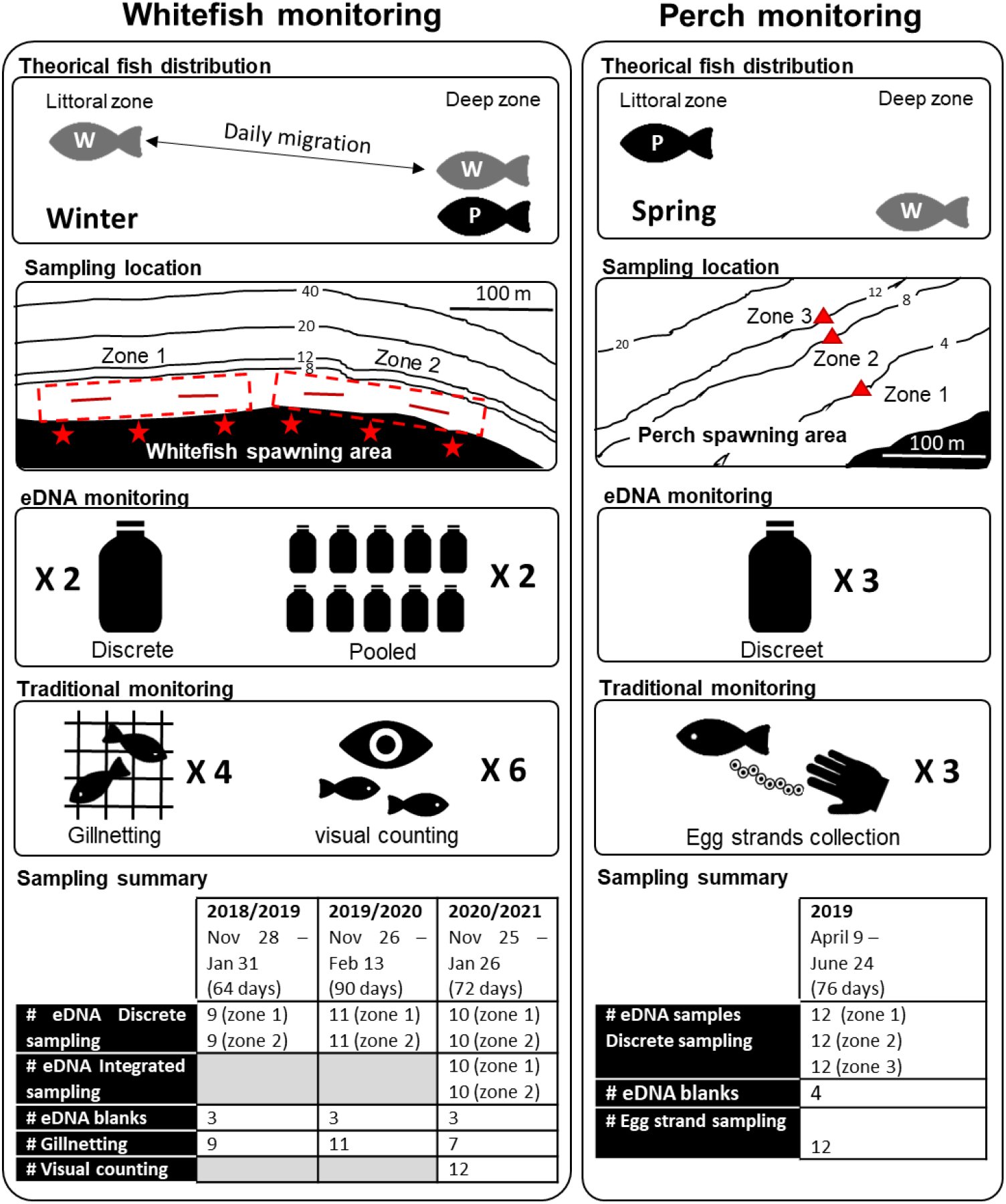
Study sites and sampling strategies in Lake Geneva. The survey was conducted during three successive winters (2018, 2019 and 2020) and one spring (2019). During winter, whitefish (W) are supposed to come to spawn in the littoral zone in the evening and to return at deeper depths in the pelagic zone during the day, while perch (P) remain in the pelagic zone. During the three years of monitoring, subsurface water samples (eDNA) were collected in the littoral zone at two stations (zone 1 and zone 2) known as whitefish spawning areas, in parallel to the traditional monitoring which consisted in the setting of 2 gillnets in each of these zones (4 red lines). In winter 2020, visual counting of whitefish present in the spawning area was performed along each of the two sampling zones with 3 observations (6 red stars), and an integrated eDNA sampling was collected in each spawning zone in addition to the eDNA discrete sampling. In spring, adult perch return to the littoral zone to spawn, while whitefish remain at deeper depths in the pelagic zone. During the spring monitoring, three artificial spawning platforms (red triangles) were set down at 4m, 8m and 12m depths to collect egg strands, and sub-surface water samples were collected each week for eDNA analysis (using discrete sampling) in the same three zones. Whitefish (W) in grey and Perch (P) in black. These maps were modified from Geodata State of Vaud, Federal Office of Topography and OpenStreetMap (https://www.geo.vd.ch/). The dates of sampling and the number of samples for both eDNA methods and traditional methods are summarized in the two tables at the bottom of the figure.

In this study, we monitored three whitefish spawning seasons during three successive winters (28^th^ Nov 2018 to 31^st^ Jan 2019 : 64 days ; 26^th^ Nov 2019 to 13^st^ Feb 2020 : 90 days ; 25^th^ Nov 2020 to 26^th^ Jan 2021 : 72 days). For perch, one year of monitoring was conducted in spring 2019 (9^th^ April 2019 to 24^th^ June 2019: 76 days); the 2020 field campaign had to be cancelled due to Covid-19 restrictions. The sampling sites are those traditionally monitored for the survey of the reproduction of these two species (Goulon *et al*., 2021). The sites are located along the beach of Ripaille in Thonon-les-bains for the whitefish monitoring (46°22’56.73’’N, 06°28’53.648’’E) and in front of the National Research Institute for Agriculture, Food and the Environment (INRAE) in Thonon-les- bains for the perch monitoring (46°22’8.663’’N, 06°27’13.946’’E) (Figure1).

### Traditional monitoring

For the survey of whitefish spawning activity, 2 benthic multimesh gillnets (L: 5 m, h = 2 m, mesh size: 19.5, 24, 29, 40, 50 et 60 mm) were set in each of the two zones of the spawning area, at a depth of about 4m. The 4 gillnets are positioned at nightfall and are pulled out the next morning. The whitefish caught in the nets are then counted and weighed. During winter 2018, 9 sampling dates were completed, 11 during winter 2019, and 8 during winter 2020 (technical reports Goulon *et al*., 2019, 2020, 2021). A minimum of 5 days and a maximum of 16 days separated two consecutive sampling dates, depending on weather conditions. In winter 2020, in addition to the gillnet monitoring, we conducted a visual counting of the number of whitefish present in the spawning area. Once or twice a week, whitefish were visually counted from the shore of the lake with the use of a torchlight, after sunset(∼7pm), during 3 minutes per site, 3 times per zone (zone 1 and 2) and always by the same expert (Fig. 1). Fish were only counted if they were formally identified by the expert. During winter monitoring 2020, 12 visual counting dates were completed; a minimum of 3 days and a maximum of 8 days separated two consecutive counting dates. The visual counting were not carried out on the same evenings as the gillnetting to avoid bias in the number of whitefish observed due to the presence of gillnets.

For the survey of perch spawning activity, three artificial spawning grounds were set down at three different depths (4, 8 and 12 m), and, once a week the artificial spawning grounds were brought out, and perch egg strands were then collected and counted (Gillet and Dubois, 1995, 2007). Twelve sampling dates were completed during spring 2019, with a minimum of 5 days and a maximum of 9 days between two consecutive sampling dates (Goulon *et al*., 2020). Unfortunately, no monitoring could be organized in 2020 due to COVID-19 restrictions.

We considered both the mean value obtained for the spawning area (2 sampling zones for whitefish and 3 zones for perch) and each value separately in order to assess the variability of the measurements within each spawning area.

### Water sampling and filtration

In total, 58 samples were collected with the discrete sampling strategy during the three winters (whitefish spawning period) with 18, 20 and 20 samples in 2018, 2019 and 2020 respectively (Fig.1). For each sample, 2 L of subsurface water were collected in each of the two zones (i.e. two samples per date). The sampling was performed before the setting of gillnets. In addition, an integrated sampling strategy was performed in 2020, allowing to collect 20 additional samples (10 for each zone). In this later case, ten 200 mL subsurface water samples were pooled to obtain the 2L to be filtered. Two pooled samples were therefore collected at each sampling date.

In spring 2019 (perch spawning period), 2L of subsurface water were collected once a week, before egg strands collection, at each of the 3 stations where the artificial spawning grounds were located. For the 12 dates of this spring survey, 36 samples were therefore collected.

All water samples were stored in cleaned bottles (washed with 10% H_2_O_2_ and rinsed 3 times with ultrapure water) immediately placed in coolers and returned to the laboratory to be filtered. Filtration was performed at the laboratory, within 2 hours after the sampling, using Sterivex ™ MILLIPORE filter units (porosity of 0.45 µm). For each filter, between 1.5 L to 2 L of water was filtered. After filtration, Sterivex were filled with preservation buffer (EDTA 40 mM, Tris-HCl (pH 8) 50 mM and sucrose 0.75 M) and stored at -20°C until DNA extraction. For all the samples, sampling and filtration were performed following the detailed protocol from Vautier *et al*. (2021), accessible at https://www.protocols.io/view/fish-edna-water-sampling-and-filtration-through-st-br5rm856.

For blank samples, cleaned bottles filled with DNA-free water were opened on the boat during sampling procedures, closed and placed in a cooler with the other samples. The blank samples were then treated in the same way as the other water samples (i.e. filtration and DNA extraction). Blank samples were performed at regular intervals during the different sampling campaigns, with a total of 13 blank samples.

### DNA extraction for water samples and fish tissues

For all the samples, DNA extraction was performed in a laboratory dedicated to the analyses of rare eDNA. Extractions were performed following the protocol from Vautier *et al*. (2020), accessible at https://www.protocols.io/view/fish-edna-dna-extraction-from-water-samples-filter-bfk8jkzw. This protocol uses the NucleoSpin® Soil kit (MACHEREY-NAGEL) with specific modifications adapted to Sterivex cartridges and buffer preservation. The DNA was eluted in 30 µL of pre-heated 55°C SE buffer, quantified with Nanodrop (Thermo Scientific) and stored at -20°C.

To test the primers’ specificity, DNA from 18 fish species (caught in Lake Bourget and Lake Geneva) was extracted from tissues with NucleoSpin® DNA RapidLyse (MACHEREY-NAGEL) (Table S1). The DNA was quantified by spectrophotometry using Nanodrop™, and stored at -20°C; these DNA extracts were used as controls to test the specificity of primer sets.

### Design of primers and probe for *Perca fluviatilis*

The specificity of primers found in the literature was tested, using PCR and qPCR assays, on DNA extracted from various fish tissues (Table S1). For the whitefish primers proposed by Hulley *et al*. (2019), we found a specific and good efficiency of amplification (Table S1); these primers were then used for ddPCR analyses to first evaluate their sensitivity and then to quantify whitefish eDNA for our environmental samples.

For the primers set proposed by Furlan and Gleeson (2016) to target perch, our tests showed that the DNA of several species present in Lake Geneva was amplified by qPCR (Table S1). We therefore designed new primers that amplified specifically perch DNA to use them for our ddPCR assays. To design these new primers, COI gene sequences were collected from Barcode of Life Database (BOLD) (http://www.boldsystems.org/) for the target species (*Perca fluviatilis)* and for 35 other fish species identified in French peri-alpine lakes. All available sequences were kept, regardless of the geographical origin, in order to account for potential intraspecific genetic variations. PCR primer-probe sequences targeting the *Perca fluviatilis* COI gene were designed using the Primers3 software (Untergasser *et al*., 2012). The detailed procedure applied to design the new primers and probe is presented in supplemental information (Text S1). The primers set selected for European perch were: Perca_Rhone_COI-5P_F (5’-CGC CGT TCT TCT CCT TCT CT- 3’), Perca_Rhone_COI-5P_R (5’-AGG AGG AGG CGA CCC TAT TC-3’) and Perca_Rhone_COI-5P_P (5’-ACT TCC TGT TCT TGC CGC TGG CA-3’) (Table 1).

**Table 1.**
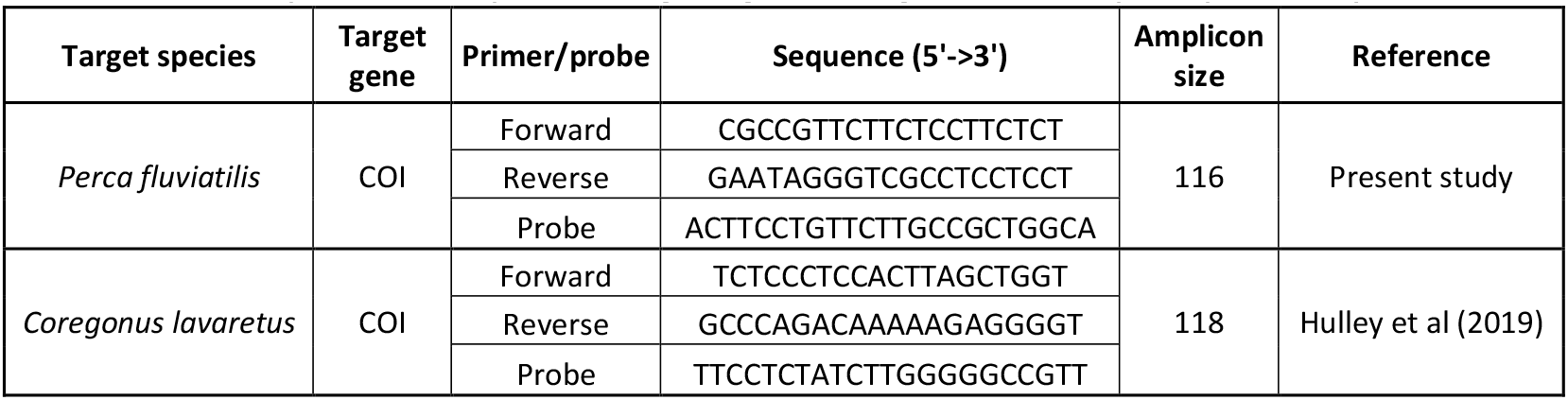
Selected primers and probes targeting the COI gene for European perch (*P. fluviatilis*) and Lake whitefish (*C. lavaretus*).

### Primers specificity and sensitivity

The specificity of primers were first tested using qPCR assays on DNA extracted from fish tissues (18 different species; Table S1). qPCR assays were performed on 100 pg of fish DNA, using the RotorGene Q (Giagen, Hilden Germany) and the Rotor-gene Multiplex PCR Kit, following the manufacturer recommendations and with a final volume of 20 µL. The qPCR program consisted of 5 min at 95 °C, followed by 40 cycles of denaturation for 10 s at 95 °C and extension at 60 °C for 40 s. qPCR data analysis was performed using the rotor-gene Q Series software (version 2.3.1).

Primers specificity was also tested in ddPCR assays with DNA pools containing DNA from 18 fish species, 17 fish species present in Lake Geneva and either perch or whitefish DNA. The DNA pools contained different amounts of fish DNA: 50, 10, 2, 0.4 and 0.08 pg. The DNA pools containing the highest amounts of DNA were processed first, and then successive dilutions by a factor 5 were performed to obtain lower concentration pools. The series included 2 replicates for each dilution and 2 negative controls. The DNA concentration for the starting dilution (50 pg) measured with ddPCR, was 89.6 copie.µL^-1^ (S.D. = 2.2) for *C. lavaretus* and 74.2 copie.µL^- 1^ (S.D. = 1.0) for *P. fluviatilis*.

To determine the sensitivity of the ddPCR assays, we conducted a serial dilution experiment with DNA extracted from tissues of *C. lavaretus* and *P. fluviatilis* to determine the limit of detection (LOD) and the limit of quantification (LOQ). The serial dilution, detailed in supplemental material (Text S2), included 5 replicates of each dilution and 5 negative controls. The starting dilution of these series (1/100 or 50 pg), measured with ddPCR, was 89.1 copie.µL^-1^ (S.D. = 4.2) for *C. lavaretus* and 74.5 copie.µL^-1^ (S.D. = 2.0) for *P. fluviatilis*, and this starting dilution was also used as a positive control in the following ddPCR analyses. The LOD was determined as the lowest concentration at which at least one positive detection was measured among the 5 replicates. The LOQ, i.e. the lowest PCR copy number concentration for which the method provides results with an acceptable uncertainty, was defined following the recommendations from Deprez *et al*. (2016) (Text S2).

### ddPCR assays applied to the environmental samples (monitoring of spawning)

ddPCR targeting *C. lavaretus* were run using the whitefish primers Forward 5’-TCT CCC TCC ACT TAG CTG GT-3’ and Reverse 5’-GCC CAG ACA AAA AGA GGG GT-3’ which amplified a 118 bp region of the COI gene and the probe 5’-TTC CTC TAT CTT GGG GGC CGT T-3’ (Hulley *et al*., 2019). To target *P. fluviatilis*, we used the primers and probe designed in this study, which amplified a 116 bp region of the COI gene (Table 2). Fluorescent dyes chosen for the hydrolysis probes were FAM (∼517nm) for *C. lavaretus* and HEX (∼556nm) for *P. fluviatilis*. ddPCR were run in multiplex reactions with both primer-probe sets multiplexed together in each PCR well (96 well plates).

**Table 2.** Minimum, maximum and mean values for perch and whitefish eDNA and traditional monitoring surveys.

|  |  |  | Winter 2018 | Spring 2019 | Winter 2019 | Winter 2020 | Winter 20 Supp. |
| --- | --- | --- | --- | --- | --- | --- | --- |
| Whitefish | Traditional<br>(Number of whitefish) | Tot | 19 (15.1 kg) | - | 13 (9.1 kg) | 5 (3.2 kg) | 397 |
|  |  | Max | 5 (4.5 kg) | - | 5 (3.2 kg) | 4 (2.5 kg) | 110 |
|  |  | Min | 0 | - | 0 | 0 | 0 |
|  | eDNA (copy.L<sup>-1</sup>) | Mean | 5 285.6 | 39.9 | 908.2 | 1 683.6 | 3 663.9 |
|  |  | Max | 27 960.0 | 178.6 | 5 013.4 | 8 491.6 | 15 126.2 |
|  |  | Min | 0.0 | 0.0 | 10.5 | 5.3 | 73.1 |
| Perch | Traditional<br>(egg strand number) | Tot | - | 281 | - | - | - |
|  |  | Max | - | 100 | - | - | - |
|  |  | Min | - | 0 | - | - | - |
|  | eDNA (copy.L<sup>-1</sup>) | Mean | 106.6 | 620.0 | 35.1 | 43.2 | 72.3 |
|  |  | Max | 307.3 | 3 505.6 | 240.7 | 89.9 | 354.9 |
|  |  | Min | 0.0 | 15.1 | 10.6 | 15.9 | 7.6 |

The eDNA extracts obtained for each sampling date were analyzed independently (*i*.*e*., 3 samples separately or 2 samples separately for spring and winter monitoring respectively, see Fig.1) and were also pooled (3 samples pooled for spring monitoring and 2 for winter monitoring) to measure an average signal for the spawning area.

The negative controls consisted of DNA-free water, whereas the positive controls consisted of DNA extracted from *P. fluviatilis* and *C. lavaretus* tissue material.

Digital droplet PCR was conducted using the Bio-Rad QX200 ddPCR system (Bio-Rad, Temse, Belgium) in a 20 μL total volume. Each reaction contained 1x Bio-Rad ddPCR supermix for probes (no dUTP), 900 nM of each primer, 250 nM probe, 3 µL of template DNA for perch monitoring and 4µl of template DNA for whitefish monitoring, from 5 U to 10 U of AflII restriction enzyme according to DNA concentration, completed with diethylpyrocarbonate (DEPC) water (Sigma-Aldrich, Overijse, Belgium). Twenty microliters of the PCR mix were pipetted into the sample chambers of a Droplet Generator DG8 Cartridge (Bio-Rad, cat no. 1864008), and 70 μL of the Droplet Generation Oil for Probes (Bio-Rad, Cat No. 186-4005) was added to the appropriate wells. The cartridges were covered with DG8 Gaskets (Bio-Rad, cat no. 1863009) and placed in a QX200 Droplet Generator (Bio-Rad) to generate the droplets. The droplets (40μL) were then carefully transferred to a ddPCR 96-well plate (Bio-Rad, Cat No. 12001925) and sealed with pierceable foil (Bio-Rad, Cat No. 181-4040). The ddPCR 96-well plates were brought into a TProfessional Basic Thermocycler from Biometra Ltd. PCR conditions were 10 min at 95 °C, followed by 40 cycles of denaturation for 30 s at 94 °C and extension at 60 °C for 1 min, with a ramp rate of 2 °C s-1, followed by 10 min at 98 °C and a hold at 4°C.

Droplets were then read on a QX200 droplet reader (Bio-Rad). All droplets were checked for fluorescence with the Bio-Rad’s QuantaSoft software version 1.7.4.0917. The fluorescence amplitude threshold, distinguishing the positive from the negative droplets was set manually by the analyst as the midpoint between the average fluorescence amplitude of the positive and negative droplet cluster. The same threshold was applied to all the wells of a given PCR plate. The average number of accepted droplets was around 17 000.

To estimate eDNA concentrations, expressed as a copy number par liter of water, calculation was performed according to the following formula, with a droplet volume (V_d_) set at 0.834 nL (Corbisier *et al*., 2015). Where C_X_ is the number of target eDNA copies per litre of 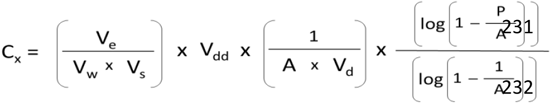 water, V_e_ is the extraction elution volume, V_s_ is the volume of DNA extract used in the ddPCR reaction, V_w_ is the volume of filtered water, Vdd is the volume of ddPCR reaction (20 µL), A is the number of accepted droplets, Vd is the droplet volume and P is the number of positives droplets.

## Results

### Validation of ddPCR assays for perch and whitefish primer sets

#### Sensitivity assays (Fig. 2)

The tests based on dilution series showed that the decline in measured concentrations followed a nearly perfect linear relation for both fish species, with a correlation coefficient r = 0.998 (p-value < 0.001) for whitefish and r = 0.999 (p- value < 0.001) for perch. The highest dilution level (1/312 500), with a fish DNA quantity of 0.016 pg, did not result in any amplification for any of the five replicates whatever the fish species. The LOD (*i*.*e*., lowest PCR copy number concentration at which at least one positive detection is measured among the 5 replicates) was 0.24 copies μl^−1^ (4.80 copies per ddPCR reaction) for whitefish and 0.11 copies μl^−1^ (2.20 copies per ddPCR reaction) for perch, and was obtained for the same dilution (1/62 500) or fish DNA quantity (0.08 pg). The LOQ (*i*.*e*., lowest PCR copy number concentration for which the method provides results with 30 % of uncertainty) was 0.63 copies μl^−1^ (12.60 copies per ddPCR reaction) for whitefish and 0.58 copies μl^−1^ (16.60 copies per ddPCR reaction) for perch, and was obtained for the same dilution (1/12 500) or fish DNA quantity (0.4 pg).

**Figure 2.**
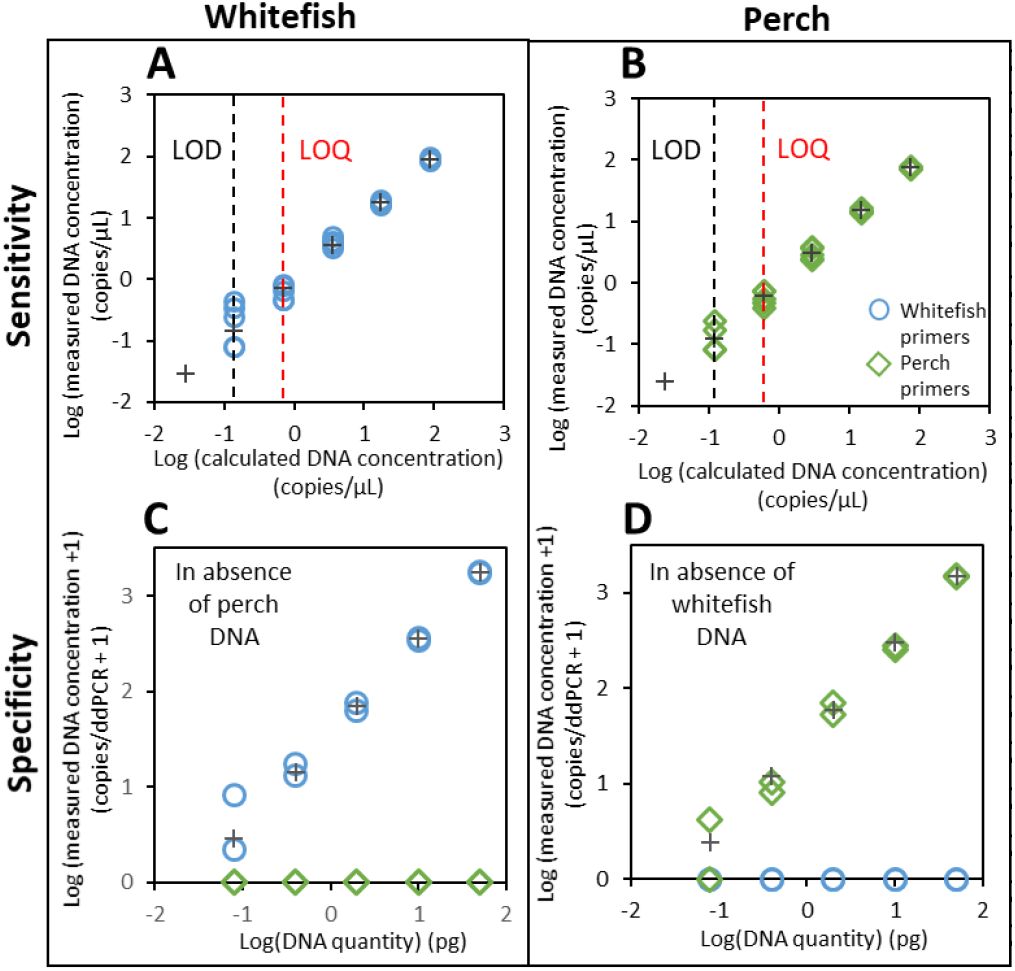
Sensitivity, limit of detection (LOD), quantification (LOQ) and specificity of the primer sets. Primers sets sensitivity and limits of detection and quantification were evaluated using six DNA quantities (50, 10, 0.4, 0.08 and 0.016 pg) obtained from five successive dilutions from the highest DNA quantity (50 pg), for whitefish (A) and perch (B) with 5 replicates each. Primers sets specificity was determined with DNA pool from 14 fish species (see Table S1) with whitefish and without perch (C), with perch and without whitefish (D). For each species, five DNA quantities (50, 10, 0.4, and 0.08pg) were used and obtained from five successive dilutions from the highest DNA quantity (50 pg). Grey cross = theoretical estimation of the target gene copies number per ddPCR well, based on gene copies number obtained from the highest DNA quantity. Whitefish primers (blue circle), perch primers (green rhomb).

#### Determination of primer-probe specificity (Fig. 2)

Whether for perch or whitefish primers selected, no non-specific amplification could be detected on the other 17 species (Table S1). When the DNA of the target species was present, the amplification was found efficient for all replicates and for all DNA amounts, except for perch for which one of the two replicates failed to be amplified with the lowest DNA amount (*i*.*e*., 0.08 pg, corresponding to the LOD). Despite the presence of DNA from other fish species, the dilution series showed that the decline in measured concentrations followed a nearly perfect linear relation for both fish species (r = 0.986 with a p- value <0.001 for whitefish and r = 0.987 with a p-value <0.001 for perch).

### DNA concentrations

For the three winter monitoring, all the 78 eDNA samples were successfully extracted. The mean DNA concentration obtained was 43.6 ± 17.9 ng μl^−1^. When two different strategies of sampling were performed in 2020, similar DNA concentrations were obtained for the discrete sampling strategy (mean 40.4 ± SD 16.7 ng μl−1) and the integrated sampling strategy (mean 39.3 ± SD 19.9 ng μl−1), with no statistical difference between the two sampling methods (T-test: p<0.05). For spring monitoring, the mean DNA concentration obtained for the 36 samples was 70.7 ± 35.5 ng μl^−1^.

In total, 13 field blank samples (controls) were collected. These blanks were found to contain low (but detectable) amounts of DNA, with an average concentration of 1.8 ± 1.1 ng μl^−1^. No amplification of fish DNA (perch or whitefish) could be obtained from these controls.

### Timing and duration of spawning periods: comparison between ddPCR quantification and traditional methods

#### Perch spawning period monitoring (Fig. 3, Table 2)

Perch eDNA could be detected at each of the 12 sampling dates for at least one of the two ddPCR replicates. The temporal dynamics are similar for both eDNA and traditional methods. Perch eDNA signal and egg strands number both increased from mid April, reach a peak early May, and decrease until early June. The two methods allowed to conclude to an equivalent spawning duration of about 8 weeks. The maximum values for perch eDNA signal and egg strands number were reached at close dates, with respectively 3 505.6 copy.L^-1^ on May 10, and 100 egg strands collected on May 2. Considering the cumulative counts, for eDNA, 50% of the perch signal was reached on May 7, consistently to what was observed for traditional monitoring (May 3). During this period, the whitefish eDNA signal remained low, and was not detected for the last two dates, in any of the two replicates. The whitefish eDNA signals showed much less variations over the time (min. 0.0 copy.L^-1^ and max. 178.6 copy.L^- 1^) in comparison to those observed forperch (min. 15.1 copy.L^-1^ and max. 3505.6 copy.L^-1^); the magnitude of the signal was approximately 20 times stronger for perch than it was for whitefish.

**Figure 3.**
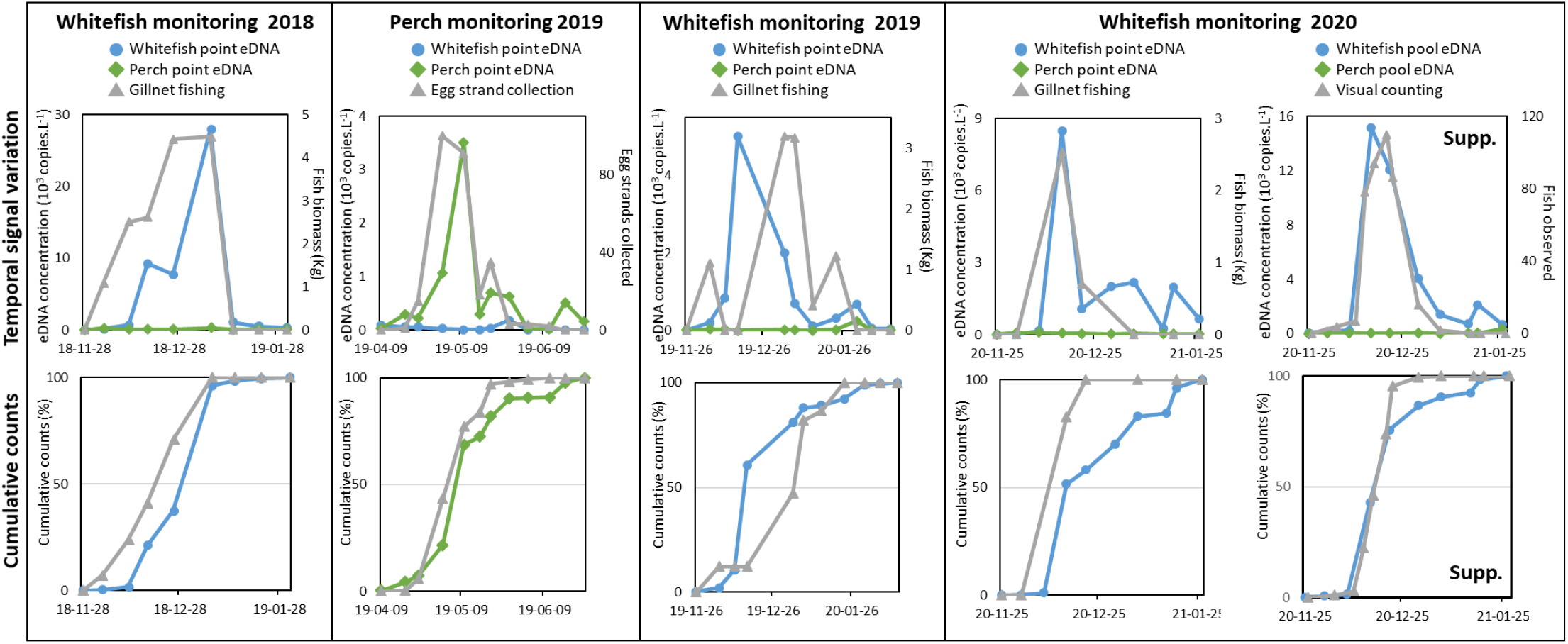
Comparison of results obtained for eDNA and traditional monitoring for the survey of the spawning phenology of whitefish (winter 2018, 2019 and 2020) and European perch (spring 2019). For whitefish, traditional monitoring consists of gillnet fishing while for perch it consists of egg strands collection. Water sampling (eDNA) was conducted using a discrete strategy (point) for all monitoring. During winter 2020, supplemental monitoring (supp.) were performed using an eDNA pooled sampling strategy and a visual counting of whitefish in the spawning areas. The results presented the average signal obtained in the spawning area (as presented previously in material and methods). The cumulative counts represent, for each date, the sum of the biomass of whitefish captured or the sum of the number of perch egg strands (traditional monitoring) or the sum of eDNA concentrations measured. Whitefish eDNA (blue circle), perch eDNA (green rhomb) and traditional monitoring (grey triangle).

#### Whitefish spawning period monitoring (Fig. 3, Table 2)

For the three years of monitoring, whitefish eDNA could be detected at each sampling date and for the two ddPCR replicates. For the three winters, both the whitefish eDNA signal and the number of whitefish caught or observed, increased from early December and decreased until mid/late January, revealing a total duration of 6 to 8 weeks according to the year. For the three monitoring periods, and for both methods (eDNA and traditional), a unique major peak was reached between mid December and early January. The peaks were concomitant for eDNA and traditional monitoring during winter 2018 (January 7) and winter 2020 (December 15), and were very close for the supplemental monitoring performed during winter 2020/21, when visual counting and integrated eDNA sampling reached a maximum on December 20 and December 15, respectively. Only during the 2019 winter, the peaks identified with the two methods were not simultaneous: the eDNA peak was reached mid December (December 16) while for traditional monitoring the maximal number of fish caught in gillnets was reached on January 3. Unfortunately no sampling could be performed between these two dates due to unfavourable weather conditions, therefore limiting our capacity to identify the peaks more precisely.

Considering the cumulative counts, for all the three eDNA surveys, 50% of the whitefish signal was reached at mid/end December (from December 14 to 28), whereas for traditional monitoring this period run on 26 days, from early December to early January (December 9 to January 4) according to the year. Over the three winters of monitoring, the perch eDNA signal could be detected at each sampling date and for at least one of the two replicates, but this signal was low and showed much less variation over the time (min. 7.6 copy.L^-1^ and max. 357.9 copy.L^-1^) than observed for the whitefish signal (min. 5.3 copy.L^-1^ and max. 27 960.0 copy.L^-1^). The magnitude of the eDNA signal was approximately 20 to 100 times stronger for whitefish than for perch.

#### Comparison of sampling methods (Fig. 3, Table 2)

Two supplemental surveys were included in the 2020 monitoring (winter): a visual counting of whitefish and an integrated eDNA sampling consisting of ten samples of 200mL pooled together to constitute the 2L of water to be filtered (in addition to the discrete sampling strategy, i.e. 2L sample collected at once). For gillnetting and visual counting, the peak was reached mid-December (December 15 and 20, respectively). However, it is noticeable that during the entire gillnet monitoring only five fish were caught, out of which four were caught at the time of the peak, whereas 397 whitefish were observed along the same shoreline, with a maximum of 110 at the time of the peak. Only seven gillnetting campaigns were performed due to bad weather conditions, while twelve visual counting campaigns were performed over the same period. A better delineation of the spawning period was therefore possible with visual counting. With regard to eDNA monitoring, for both integrated and discrete samplings, the peaks and the 50% of cumulative counts were reached mid December at very close dates (15-16 Dec) with similar temporal profiles. However, the whitefish eDNA signal was almost twice stronger with the integrated sampling (max 15126.2 copy.L^- 1^ and mean 3 663.9 copy.L^-1^) as compared to the discrete sampling (max 8491.6 copy.L^-1^ and mean 1 683.6 copy.L^-1^). Interestingly, the perch eDNA signal was also stronger with the integrated sampling (max 354.9 copy.L^-1^ and mean 72.3 copy.L^-1^) compared to the discrete sampling (max 89.9 copy.L^-1^ and mean 43.2 copy.L^-1^), but remained, as expected, at a much lower level in comparison to whitefish eDNA signal.

### Spatial variability of the eDNA signal (Fig. 4, Supplemental information Table S2.)

Here we considered independently the ddPCR results obtained for each sampling zones of the spawning areas (3 zones for the spring survey and 2 zones for winter surveys) to assess the variability of the eDNA signal between the different zones.

**Figure 4.**
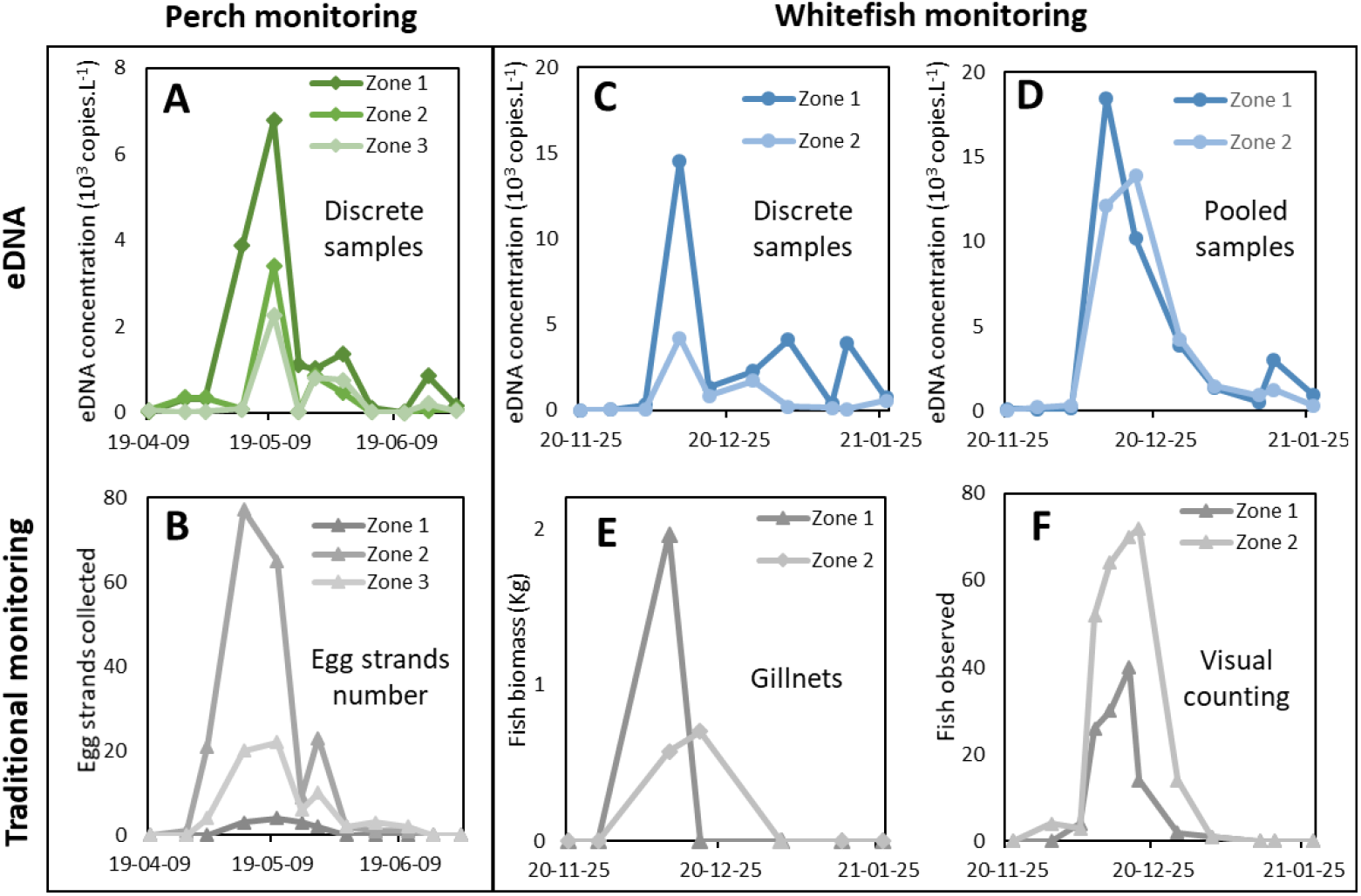
Spatial variability of eDNA signal and traditional monitoring for perch and whitefish during their respective spawning periods. For perch monitoring: variability of the perch eDNA signal illustrated by the 3 replicates obtained from discrete sampling performed in zones 1, 2, and 3 (**A**) and variability of the number of perch egg strands collected at the three artificial spawning platforms positioned respectively in zones 1, 2 and 3 (**B**). For whitefish monitoring: variability of the withefish eDNA signal illustrated by the 2 replicates obtained from discrete sampling performed in zones 1, 2 (**C**) or 2 replicates obtained from pooled water samples performed in zones 1, 2 (**D**), and whitefish biomass caught in gillnets positioned in the two spawning zones 1 and 2 (**E**) or number of whitefish observed by visual counting at the two spawning zones 1 and 2 (**F**). Whitefish eDNA (blue circle), perch eDNA (green rhomb) and traditional monitoring (grey triangle).

#### Spring monitoring

Perch eDNA and traditional monitoring were performed in three distinct zones (zones 1 and 2 are distant from 65 m, and 20 m separate zones 2 and 3). For both approaches and for the three zones, the spawning peak was reached at the beginning of May (between May 2 and 10), but the intensity of the peaks varied significantly between the zone 1 and the two other zones (Fig.4). For eDNA monitoring, the highest peak was observed in zone 1 (6 780.3 copy.L^-1^), and the lowest in zone 3 (2 270.2 copy.L^-1^). Regarding the traditional monitoring, we also observed significant variations in peaks intensity between the zone 1 and the two other zones, but the highest peak was, in this case, observed in zone 2 (77 egg strands), and the lowest in zone 1 (4 egg strands).

#### Winter monitoring

During winter 2020, whitefish spawning monitoring was performed in two distinct zones of the spawning area (each zone is about 250m long and the two zones are separated by about 30 meters), using two eDNA approaches (discrete and integrated sampling) and two traditional approaches (gillnetting and visual counting). For the four approaches and the two zones, the spawning peak was reached around mid-December (between December 15 and 22). However, the intensity of the peaks varied between the two zones, with a highest signal found in zone 1 for both eDNA monitoring and gillnetting, whereas a highest signal was detected in zone 2 for visual counting (Fig.4). For the eDNA method, the integrated sampling strategy appeared to reduce the variability of eDNA signals in comparison to the discrete sampling strategy (Fig.4, Table S2). For integrated samples the difference of intensity between the two peaks (18 489.9 copy.L^-1^ for zone 1 and 13 900.4 copy.L^-1^ for zone 2) is moderate, while for discrete samples the peak in zone 2 (4 207.0 copy.L^-1^) is much weaker than the one found in zone 1 (14 538.7 copy.L^-1^). The averaged eDNA values obtained over the whole temporal survey were similar in zone 1 and in zone 2 for integrated samples (mean 3 878.0 copy.L^-1^ and 3 483.9 copy.L^-1^, in zone 1 and 2, respectively), while an important difference between zone 1 (mean 2 765.0 copy.L^-1^) and zone 2 (mean 781.6 copy.L^-1^) was found with discrete samples. Concerning the traditional monitoring, for gillnetting, the total number of whitefish caught was very low (3 in zone 1 and 2 in zone 2), while with visual counting the number of whitefish observed was significantly higher in zone 2 (280 in total, with a peak reaching 72) than in zone 1 (117 in total, with a peak at 40).

## Discussion

### Quantification of eDNA to monitor fish spawning in lakes

We demonstrated here the applicability of an eDNA quantitative method for the survey of whitefish and perch spawning phenology in lakes. Despite the fact that whitefish and perch live and reproduce in lakes (they are non-migratory species), their respective spawning period and the associated peak of reproduction were clearly detectable using eDNA protocols based on multiplex ddPCR assays. The operational non-invasive method we propose here with eDNA to monitor fish spawning in lakes could provide useful data with direct implications for fish management. For instance, whitefish spawning time fluctuates considerably from one year to another (Hartmann, 1984) and knowing the start and the end of the reproduction period is essential to adapt the closing dates of the fishing period. Other potential applications are also to be explored, as for instance the capacity to approximate the number of mature perch female during spawning, since we showed a good concordance between eDNA signal and the number of egg strands collected and we know that the number of egg strands reflects the number of mature females present (Gillet, Lang and Dubois, 2013).

To quantify fish eDNA, we chose to use ddPCR. In comparison to qPCR, digital PCR is known to be more reproducible (Bipm, 2012), more precise (Sanders *et al*., 2011), faster and cheaper (Nathan *et al*., 2014), and more tolerant to the presence of PCR inhibitors (Doi, Takahara, *et al*., 2015; Hoshino and Inagaki, 2012). The use of ddPCR is currently increasing in various types of environmental surveys of fish or other species in freshwaters, coastal and marine ecosystems (Crane *et al*. 2020). The efficiency and specificity of primers used in such PCR-based assays is obviously of great importance. As recommended by Goldberg *et al*., 2016, we performed *in silico* and *in vitro* testing to ensure that our primers specifically target the expected species in the studied environment, both for the new primers designed for perch and the already available ones for whitefish (Hulley *et al*., 2019). ddPCR is known to provide precise measurement without the need of calibration (White *et al*., 2009), we however performed calibrations, as recommended by the digital Minimum Information for Publication of Digital PCR Experiments (MIQE) guidelines (Huggett *et al*., 2013), in order to allow a robust comparability of the measurements obtained for perch and whitefish. LOD and LOQ calculation allowed to determine that the whitefish primers set, originally designated for qPCR, were effective in ddPCR, as is the perch primers set designed in this study, with similar efficiency for the two sets of primers.

The weekly eDNA monitoring allowed to estimate the beginning and the end of the spawning period for both species, perch and whitefish. We observed a good consistency between eDNA results and the observations made by traditional methods, with concomitant, or nearly concomitant peaks. The few-days time lag between the whitefish eDNA peak and the visual counting peak (year 2020) is explained by the fact that the counting were not performed exactly the same days as eDNA samplings. As for the differences observed between the perch eDNA peak, detected a bit more earlier than the maximum perch egg strands number, we assume that the measured eDNA signal is not only related to egg strands numbers but also to the intense activity of fish gathering into the spawning area before the eggs strand were all released.

Overall a good match is therefore observed between traditional and eDNA methods. The only exception was in winter 2019, when an important time-lag (18 days) was found between the eDNA peak and the one found with gillnetting, which could be easily explained by the fact that some of the gillnet samplings were missing (due to bad weather conditions) exactly at the time of the expected spawning peak (end of December). Additionally, the quantitative data provided by gillnet sampling were not considered robust due to the relatively low total number of fish caught (a total of 5 fish in 2020, 13 in 2019 and 19 in 2018) while, with 397 observed whitefish, visual counting seems more robust for the comparison with eDNA.

#### Use of an in situ eDNA control

Perch and whitefish spawning occur at different seasons, making it possible to use one species as a “control” for the other one. We systematically quantified the eDNA signal of the two fish species for each sampling dates, to distinguish the basal eDNA signal emitted by a fish species that lives around or in the studied zone, from the signal associated to the actual spawning activity. The increase in eDNA signal during the spawning season has been reported in several studies (e.g. Laramie, Pilliod and Goldberg, 2015; Erickson *et al*., 2016; Stewart *et al*., 2017); however, to our knowledge, this is the first time that a control species has been used to define an eDNA basal signal in order to confirm that the increase of the eDNA signal of the spawning species is not an artifact due to a phenomenon that affects the whole eDNA signal (e.g. MES, pH, wind, water flows, or methodological biases during sampling, filtration or eDNA extraction). We found that the amplitude of the eDNA signal was between 20 and 100 times higher for the spawning fish species than for the ‘control species’. The basal signal of both species has been found to be quite similar and low for both species (mean 39.9 copies per L for whitefish and 66.6 copies per L for perch), while it increased up to 27 960.0 copies per L for whitefish and 3505.6 copies per L for perch during their respective spawning activity peak. Basal eDNA signal have been detected for each sample, except at the end of June 2019 when whitefish eDNA signal was not detectable. Because whitefish live in depth during summer to avoid warm temperatures, we assume that its eDNA signal was more difficult to capture in the sub-surface water when the lake was stratified, as it was demonstrated for the eDNA signal of deep-water species in other lakes (Hänfling *et al*., 2016; Littlefair *et al*., 2021). It is therefore important to choose carefully the “control species” in order to maximise the chance of getting a sufficiently stable eDNA signal.

#### Localisation and persistence of eDNA signal in water

Poorly studied in lakes, but generally considered in mesocosm experiments (e.g. Sassoubre *et al*., 2016; Lacoursière-Roussel, Rosabal and Bernatchez, 2016), the persistence of the fish eDNA signal in lake water is a key question for the interpretation of *in situ* eDNA signal. Here, we observed a marked decrease in the quantity of eDNA signal between two successive weekly sampling at the end of the spawning periods, which suggest that the eDNA signal does not persist longer than one week. For the monitoring of spawning activity, the rather short persistence time of eDNA in aquatic environment is an advantage, allowing for close, near real-time, monitoring of the fish presence. It would therefore be relevant to carry out eDNA monitoring with a shorter timescale (e.g. twice a week, every day), as it has already been done in river (e.g. (Thalinger *et al*., 2019), in order to get a more precise timing of the reproductive activity in lakes.

From our data, we estimated that the eDNA signal however persisted in water for at least a dozen hours close to the location where he was emitted. We indeed found high amount of eDNA in the water collected in the late afternoon, i.e. approximately 12 hours after whitefish had left the spawning area for its nycthemeral migration (Eckmann, 2011; Gillet, 1989). Even if the spawning activity occurs at night, the eDNA signal can therefore be captured during the following day in lakes, this could ease the monitoring compared to surveys that have to be conducted after nightfall, such as visual counts.

Our observations on eDNA persistence in the littoral zone of Geneva Lake are however to be extrapolated to other lakes with certain precautions since the persistence, degradation and transport of eDNA signals in water are influenced by many factors, such as eDNA characteristics, abiotic environmental factors and biotic factors (Harrison, Sunday and Rogers, 2019; Wang *et al*., 2021). These complex mechanisms vary according to the species and environments studied, and divergent results have been obtained regarding eDNA degradation in freshwaters (Mauvisseau *et al*., 2021), improving our understanding of eDNA persistence in the aquatic environment is clearly needed to help interpret future eDNA surveillance results.

#### Spatial variability of eDNA signal

The eDNA transport, widely studied in lotic systems as rivers where eDNA can be transported from meters to kilometers (Pont *et al*., 2018; Jane *et al*., 2015; Deiner *et al*., 2016), is theoretically less problematic in lentic waters. However, the vertical stratification/mixing cycles are known to influence the distribution of eDNA (Lawson Handley *et al*., 2019; Littlefair *et al*., 2021), and more globally, all the characteristics of the lake hydrodynamics have to be taken into account. The effects of horizontal water movements on the distribution of eDNA in lakes have poorly been studied, but in our case study we assume that horizontal transport partly explained the local spatial variability of eDNA. It has been shown that fish eDNA in water can be assimilated to particles that can therefore be transported (Sassoudre *et al*., 2016 ; Turner *et al*., 2014). The particle tracker developed by EPFL for Geneva lake (http://meteolakes.ch/#!/hydro/geneva) allowed us to observe that, during the whitefish monitoring 2020, the lake surface particles followed a north-south direction and could therefore explain why we obtained a strong eDNA signal in the southern zone (zone 1) while a higher number of whitefish were observed in the north part of the spawning area (zone 2).

Whatever the eDNA sampling strategy we applied (discrete or integrated water samples), we were able to identify the fish spawning periods and the peak of reproduction. However, with discrete sampling strategy, we observed a spatial variability in the eDNA signal, i.e. variability for samples collected only ten meters apart within the spawning area, for both perch and whitefish monitoring. It has already been shown that eDNA is patchily distributed in ponds or lakes and that the eDNA signal decline over short distances between few to hundreds of metres (Pilliod, Goldberg, Arkle, & Waits, 2013; Eichmiller, Bajer, & Sorensen, 2014Lawson Handley *et al*., 2019). However, here we highlight that even during fish spawning activity, while huge eDNA amounts are released, the signal is not uniformly distributed over the entire spawning area, and that there is a risk to under- or over-estimate the eDNA spawning signal in particular if the sampling strategy is based on one single discrete sample. The exact location where the highest number of egg strands was observed was not necessarily the location where the highest perch eDNA quantity was found with discrete samples. Sampling is a crucial step for eDNA approaches, we point out here the need to take into account the potential eDNA spatial variability within the spawning area. An integrated sampling strategy can help smoothing the potential heterogeneity of eDNA signal, for instance by using the pooling of multiple samples (or possibly other methods to collect integrative sampling; e.g. sampling used by Civade *et al*. (2016)). We showed that pooling multiple sub-samples from the same sampling station indeed allowed to obtain very constant timing for the peaks of spawning activity in the two monitored zones (also consistent with traditional monitoring), in addition it allowed to obtain a stronger eDNA signal in comparison to the discrete strategy. Even if it has been shown that pooling multiple samples decreases the probability to detect rare eDNA signals (Sato *et al*., 2017b), it can reduce the variability of the high eDNA signal released during spawning activity. Consequently, we would recommend using the integrated sampling approach to monitor the eDNA released during fish spawning.

#### Inter-annual variability of the whitefish eDNA signal

The intensity of the eDNA signal (peaks and average) found for whitefish varied over the three years of winter monitoring (max. from 5013.4 copy.L^-1^ to 27960.0 copy.L^-1^, and mean from 908.2 copy.L^-1^ to 5293.7 copy.L^-1^). The ecological interpretation of this inter-annual differences and the comparison of the eDNA amounts measured during the spawning peaks should be taken with caution due to the frequency of sampling (weekly) and to the fact that some sampling campaigns missed in December. It has been established that fish biomass is correlated with the amount of eDNA released (Takahara *et al*., 2012). This inter-annual fluctuations could therefore reflect changes in the abundance of whitefish in the spawning area, and/or could be correlated to the number of oocytes released into the environment (also linked to the number of females), but could also be explained by differences in environmental conditions (e.g. MES, pH, hydrodynamics, etc) from one year to another. From our data, we cannot conclude about the relative importance of these potential explanatory factors. However, because the perch eDNA signal followed the same trend (though in lower quantities) as whitefish, with lower average concentrations measured in 2019 for the two species, we assume that the entire eDNA fish signal was affected the same way and, in consequence, that the interannual variability was partly driven by changes in environmental conditions.

## Conclusion and perspectives

Here, we provided a quantitative method that is applicable for eDNA survey of whitefish and perch spawning phenology in lakes, with new primers designed to target specifically *P. fluviatilis* and optimized for ddPCR use. The eDNA spawning monitoring method developed here was shown to be effective to identify the beginning, the end and the peak of the spawning period of the two targeted species. To be implemented in the future as a monitoring tool, we point out the need to carefully choose the sampling site, preferably based on existing knowledge regarding spawning area, unless the objective of the study is to explore potential –unknown- spawning areas. Additionally, an integrated sampling method to collect water would be recommended, as we observed a spatial variability of the eDNA signal within the studied spawning areas. Also, to maximize detection, DNA isolated from all states (e.g. intracellular, dissolved, particle absorbed) could be utilized, by performing water filtration using different pore sizes as recommended by (Mauvisseau *et al*., 2021). The persistence, degradation and transport of eDNA signals in lakes are complex mechanisms involving multiple biotic and abiotic factors, our understanding of these mechanisms is still limited and further study of how environmental parameters affect eDNA decay in aquatic environments is needed.

Although our eDNA method is effective to characterise the spawning activity of perch and whitefish in lakes, optimisations could still be gained by exploring the use of other marker genes. Here, we targeted mitochondrial DNA, but nuclear DNA could be targeted to measure the fish spermatozoa eDNA signal as previously shown by Bylemans *et al*. (2017) in rivers. In addition, as spermatozoa are generally distributed more homogeneously within the waterbody (Cosson *et al*. 2008), targeting nuclear DNA may allow reducing the spatial variability of the eDNA signal observed with mitochondrial DNA.

Another potential improvement for the monitoring of fish spawning is the use of eRNA that has been shown to be as sensitive as eDNA, but with a better correlation to fish abundance (Miyata *et al*. 2021). eRNA should therefore allow a better estimation of the number of fish that spawn or the quantity of gametes released. As eRNA has a shorter lifetime than eDNA in the environment (Marshall *et al*. 2021), the use of the eRNA/eDNA ratio could also be used to improve the interpretation of detections for spawning activity. The highest eRNA/eDNA ratios should reveal a signal that is freshly released in the environment, allowing to reduce potential error due to legacy DNA signal.

Though eDNA monitoring is a very useful complement to traditional methods, allowing to reduce the negative impact or failures/limitations of some traditional approaches as gillnetting, some information are however obviously not obtained via eDNA (e.g. size, age, sex of fishes). The two types of methods are therefore to be used in a complementary way, for instance using gillnets only at the time of the reproduction peak to obtain information on life traits on the fish.

## Acknowledgements

We thank the Observatory on Lakes (OLA) for the services this infrastructure has provided in particular for the collection of data based on traditional monitoring of fish spawning phenology. J.C. Hustache, P. Perney, L. Espinat provided help for field campaigns (artificial spawning ground to collect perch egg strands). We wish to thank CIPEL (Commission Internationale pour la Protection des Eaux du Léman) who indirectly supported this study through the long-term monitoring of Geneva Lake (Léman).

## Funding

This work was supported by the French Office for Biodiversity (OFB) France

## SUPPLEMENTAL INFORMATION

### Text S1. Supplemental methods - Design of primers targeting European Perch

The following parameters were applied to design forward and reverse primer pairs: minimum and maximum PCR product size of 60 to 160 respectively, minimum and maximum primer melting temperature of 50 to 65°C with a maximum temperature difference of 3°C between forward and reverse primers. As well, corresponding probe parameters included minimum and maximum size of 18 and 27 nucleotides respectively, a minimum and maximum primer melting temperature of 60 and 67°C and a GC content between 30% and 80%.

For the perch primer sets generated, specificity was determined using COI gene sequences from all the other 35 fish species selected previously. All the sequences were aligned using a multiple sequence alignment software, MEGA7 (Kumar, Stecher and Tamura, 2016). For each of the *Perca fluviatilis* primer sets (forward, reverse and probe), only primer sets with no perfect match with any of the 35 other fish species sequences were selected. Due to the high degree of homology between some fish species, COI sequences for all the 35 fish species were aligned against the selected primer and probe sequences, and the number of base pair mismatches were summed to select only primer sets with the most mismatches. The secondary structure of the selected amplicon was then checked using the Mfold program (Zuker, 2003), and finally, the amplicon was blasted in NCBI database (NCBI, 2016) to verify it only targets the *Perca fluviatilis* genome.

### Text S2. Supplemental methods – Evaluation of primers sensitivity in ddPCR assays

To determine the relative sensitivity of the ddPCR assays, we conducted a serial dilution experiment with positive controls (DNA extracted from tissues of *C. lavaretus* and *P. fluviatilis*) to determine the limit of detection (LOD) and the limit of quantification (LOQ). The serial dilution was run with a total cellular DNA sample concentrated at 1 ng μl−1, that was first 1:100 diluted to obtain a starting point for a five-step series of 5-fold dilutions (1/100, 1/500, 1/2500, 1/12 500, 1/ 62 500 and 1/312 500), and therefore six different DNA quantities (50, 10, 2, 0.4, 0.08 and 0.016 pg) according to the recommendations of Brys *et al*. (2020). The series included 5 replicates of each dilution and 5 negative controls (DNA free water instead of fish DNA). The starting dilution of these series (1/100 or 50 pg), measured with ddPCR, was 89.1 copie.µL^-1^ (S.D. = 4.2) for *C. lavaretus* and 74.5 copie.µL^-1^ (S.D. = 2.0) for *P. fluviatilis*. The LOD is defined as the lowest PCR copy number concentration that can be reliably distinguished from the negative controls (Biorad, 2016; Kiselinova *et al*., 2014), and as no amplification was obtained from any negative control, LOD was determined as the lowest concentration at which at least one positive detection was measured among the 5 replicates. The LOQ is defined as the lowest PCR copy number concentration for which the method provides results with an acceptable uncertainty. We choose an uncertainty of 30 % for a measurement result obtained as an average of the five replicate measurements, following the recommendations of Deprez *et al*., 2016.

**Table S1:** Fish species used to test the specificity of perch (*Perca fluviatilis*) and whitefish (*Coregonus lavaretus*) primers sets, and to develop new primers to target perch. For each fish species the number of distinct nucleotides with the primer sets is presented as the mismatches sum with the forward, the reverse and the probe. The specificity of primers was also tested using qPCR assays against DNA extracted from tissue of 18 fish species (*), qPCR results are expressed in cycle threshold (Ct).

| Fish species | Perch primer set<br>(Furlan and<br>Gleeson 2016) |  | Perch primer set<br>(present study) |  | Whitefish primer<br>set<br>(Hulley et al. 2019) |  |
| --- | --- | --- | --- | --- | --- | --- |
|  | Mismatches | Ct | Mismatches | Ct | Mismatches | Ct |
| *Abramis brama* * | 14 | 35.4 | 14 | - | 15 | - |
| *Alburnoides bipunctatus* | - |  | 20 |  | 18 |  |
| *Alburnus alburnus* | - |  | 13 |  | 16 |  |
| *Ameiurus melas* * | 16 | - | 19 | - | 18 | - |
| *Barbatula barbatula* * | 16 | - | 15 | - | 13 | - |
| *Barbus barbus* * | 14 | - | 16 | - | 18 | - |
| *Blicca bjoerkna* | 13 |  | 17 |  | 14 |  |
| *Coregonus lavaretus* * | 12 | - | 18 | - | 0 | 28.5 |
| *Cottus gobio* | 8 | 37.8 | 12 |  | 15 |  |
| *Cyprinus carpio* | 15 |  | 13 |  | 18 |  |
| *Esox lucius* * | 11 | 36.0 | 15 | - | 18 | - |
| *Gasterosteus aculeatus* | 8 |  | 18 |  | 19 |  |
| *Gobio gobio* * | 13 | 38.4 | 13 | - | 18 | - |
| *Gymnocephalus*<br>*cernuus* * | 1 | 36.5 | 14 | - | 15 | - |
| *Lampetra planeri* | - |  | 13 |  | 16 |  |
| *Lepomis gibbosus* * | 13 | 36.3 | 18 | - | 17 | - |
| *Leuciscus leuciscus* | - |  | 16 |  | 17 |  |
| *Lota lota* * | 16 | - | 15 | - | 17 | - |
| *Oncorhynchus mykiss* | 12 |  | 20 |  | 13 |  |
| *Perca fluviatilis* * | 0 | 28.2 | 0 | 27.7 | 14 | - |
| *Phoxinus phoxinus* | 16 |  | 18 |  | 17 |  |
| *Pseudorasbora parva* | 12 |  | 15 |  | 19 |  |
| *Rhodeus amarus* | 16 |  | 18 |  | 18 |  |
| *Rutilus rutilus* * | - | - | 14 | - | 16 | - |
| *Salaria fluviatilis* | - |  | 17 |  | 13 |  |
| *Salmo trutta* * | 13 | - | 18 | - | 7 | - |
| *Salvelinus alpinus* * | 10 | - | 18 | - | 10 | - |
| *Sander lucioperca* | 0 |  | 14 |  | 16 |  |
| *Scardinius*<br>*erythrophthalmus* * | 13 | - | 17 | - | 16 | - |
| *Silurus glanis* * | 16 | - | 18 | - | 16 | - |
| *Squalius cephalus* * | 12 | 36.9 | 16 | - | 15 | - |
| *Telestes souffia* | - |  | 16 |  | 14 |  |
| *Thymallus thymallus* | 14 |  | 15 |  | 7 |  |
| *Tinca tinca* * | 12 | 36.1 | 11 | - | 17 | - |
| *Zingel asper* | - |  | 12 |  | 17 |  |

**Table S2:** Wilcoxon test results for traditional and eDNA monitoring to compare the results obtained for different zones (within each spawning area), for perch (zones 1, 2 and 3) and for whitefish (zones 1 and 2). In grey, the comparisons for which a significant difference was found.

|  |  | Perch |  |  | Whitefish |  |
| --- | --- | --- | --- | --- | --- | --- |
|  |  | Zone 1 vs<br>zone 2 | Zone 1 vs<br>zone 3 | Zone 2 vs<br>zone 3 | Point<br>zone 1 vs<br>zone 2 | Pool<br>zone 1 vs<br>zone 2 |
| eDNA<br>METHODS | Wilcoxon | 9 | 2 | 17 | 1 | 26 |
|  | Critical value (0,05%) | 13 | 13 | 13 | 8 | 8 |
|  | P-value | 0,02 | 0,002 | 0,1 | 0,005 | >0,2 |
|  |  | Zone 1 vs<br>zone 2 | Zone 1 vs<br>zone 3 | Zone 2 vs<br>zone 3 | Nets<br>zone 1 vs<br>zone 2 | Visual<br>counts<br>zone 1 vs<br>zone 2 |
| TRADITIONAL<br>METHODS | Wilcoxon | 0 | 0 | 23 | 7 | 6 |
|  | Critical value (0,05%) | 13 | 13 | 13 | 2 | 13 |
|  | P-value | <0,001 | <0,001 | >0,2 | >0,2 | 0,007 |

## Notes

### Competing Interest Statement

The authors have declared no competing interest.

